# Impact of *Cymbopogon flexuosus* (Poaceae) essential oil and primary components on the eclosion and larval development of *Aedes aegypti* (Diptera: Culicidae)

**DOI:** 10.1101/2020.12.30.424851

**Authors:** Ruth Mariela Castillo-Morales, Sugey Ortiz Serrano, Adriana Lisseth Rodríguez Villamizar, Stelia Carolina Mendez-Sanchez, Jonny E. Duque

**Affiliations:** Centro de Investigaciones en Enfermedades Tropicales - CINTROP. Facultad de Salud. Escuela de Medicina, Departamento de Ciencias Básicas, Universidad Industrial de Santander, Guatiguará Technology and Research Park, Km 2 Vía El Refugio, Piedecuesta, Santander, Colombia.; Grupo de Investigación en Bioquímica y Microbiología (GIBIM). Escuela de Química, Universidad Industrial de Santander, Bucaramanga A.A. 678, Colombia

**Keywords:** Juvenile hormone, molt hormone, methoprene, diflubenzuron

## Abstract

The current study describes the effects of sub-lethal concentrations and constituent compounds (citral and geranyl acetate) of *Cymbopogon flexuosus* essential oil (EO) on the development of *Aedes aegypti* (L.) eggs and larvae. To demonstrate the ovicidal activity of EO, we treated embryonated eggs with 6, 18, and 30 mg.L^-1^ and larvae with 3 and 6 mg.L^-1^ EO concentrations. Citral and geranyl acetate were evaluated at 18, 30, and 42 mg.L^-1^ and compared with commercial growth inhibitors (diflubenzuron and methoprene) at 3 and 6 mg.L^-1^ concentrations. For each treatment, we measured larval head diameter, siphon length, and body length. To complement the morphological analysis, we examined concentrations of moult hormone (MH) and juvenile hormone III (JH III) using high-performance liquid chromatography (HPLC) coupled to mass spectrometry. The EO decreased egg hatching at all concentrations: 6 mg.L^-1^ in 45.3%; 18 mg.L^-1^ in 23.3%, and 30 mg.L^-1^ in 34.6%. EO also altered molting among larval instars and between larvae and pupae, with an increase in the length (3 mg.L^-1^: 6 ± 0.0 mm; 6 mg.L^-1^: 6 ± 0.7 mm) and head width (3 mg.L^-1^: 0.8 ± 0 mm; 6 mg.L^-1^: 0.8 ± 0.0 mm) compared with the control group (length: 5.3 ± 0 mm; head width: 0.7 ± 0.0 mm). We did not detect chromatographic signals of MH and JH III in larvae treated with *C. flexuosus* EO or their major compounds. The sub-lethal concentrations *C. flexuosus* EO caused a similar effect to diflubenzuron, decreasing hormone concentration, extending the larval period, and death.

## Introduction

Annually, more than 2.5 billion people are a risk to get different arboviruses like Dengue, Zika, and Chikungunya virus in urban and peri-urban areas due to the vector *Aedes aegypti* (Guzman *et al*. 2010, Bhatt *et al*. 2013). Given that there is no effective vaccine for any of these three diseases (Laughlin *et al*. 2012, Rather *et al*. 2017), the principal strategy to mosquitoes control consists in the population decrease using chemical insecticides on juvenile stages (temephos) and adults (deltamethrin, lambda-cyhalothrin, cyfluthrin, and malathion) (Baldacchino *et al*. 2015, Rather *et al*. 2017). However, the inappropriate use and application of these products generate resistant populations to the active principles, environmental deterioration, and non-target organisms’ effects (Harris *et al*. 2010, Santacoloma Varón *et al*. 2010, Lima *et al*. 2011, Ardila-Roldán *et al*. 2013, Grisales *et al*. 2013).

In the search for new mosquito control alternatives, the juvenile stages are underexplored as an ideal target of control because these stages have developed exclusively in aquatic environments. Any interferences or alterations in these environments caused by an external source of hormonal or growth regulation can interrupt the normal life cycle and cause abnormal development in adult stages (Hoffmann and Lorenz 1998). According to this, one of the control strategies with major specificity and minor probability to generate resistance are the insect growth regulators (IGR), these differ from the conventional insecticides because they cause morphological and physiological changes during the development and metamorphosis (Baldacchino *et al*. 2015, Rather *et al*. 2017). The most used and studied growth regulators are diflubenzuron (inhibitor of chitin synthesis) and methoprene (juvenile hormone analogue) (Licastro *et al*. 2010, McGraw and O’Neill 2013).

Moving forward in search of more products to control mosquitoes, different scientific studies with essential oils (EO) have reported repellent, deterrent, ovicidal, larvicidal, pupicidal, and adulticidal activities against *Ae. aegypti* (Ponnusamy *et al*. 2010, Aciole *et al*. 2011, Baldeón 2011, Govindarajan *et al*. 2011, El-Gendy and Shaalan 2012, Elango *et al*. 2012, Gokulakrishnan *et al*. 2013, Suman *et al*. 2013, Vera *et al*. 2014, Castillo *et al*. 2017). These studies mentioned that EO from the Malvaceae, Asteraceae, Annonaceae, Rutaceae, Piperaceae, and Verbenaceae plant families are metamorphosis inhibitors that caused reproductive damages in females (Shaalan *et al*. 2005, Rafael *et al*. 2008, Arivoli and Tennyson 2011, Dua *et al*. 2013). Although ovicidal and larvicidal activity was reported in plants of genera *Cymbopogon* against *Ae. aegypti* (Freitas *et al*. 2010, Warikoo *et al*. 2011), Soonwera and Phasomkusolsil (2016) reported morphological abnormalities over juvenile stages of *Ae. aegypti* and *Anopheles dirus* larvae and deformed pupae, as well as incomplete hatching and mortality due to the action of *Cymbopogon citratus* EO.

Previously, studies evaluated the activity of *Cymbopogon flexuosus* (Poaceae) EO against *Ae. aegypti* in terms of larvicidal activity, repellence, or dissuasive efficacy using a lethal concentration (LC) (CL50 = 17.2 mg.L^-1^; CL95 = 49.9 mg.L^-1^) (Vera *et al*. 2014, Castillo *et al*. 2017). However, these studies did not mention the effect of this EO on development alterations. Therefore, the objectives of this study were to describe the effect on *Ae. aegypti* eggs and larval development when treated with of sub-lethal concentrations of *Cymbopogon flexuosus* EO and its major compounds: citral (consist of a mixture of two isomers geranial and neral) and geranyl acetate. Additionally, considering that the major compounds of this plant have structural similarity (Figure 1) with the commercial growth regulators and developmental inhibitors, we used high-performance liquid chromatography (HPLC) coupled with mass spectrometry to examine the potential variations of the concentrations of moult hormone (MH) and juvenile hormone III (JH III) caused by *C. flexuosus* EO and its major compounds on *Ae. aegypti* larvae.

**Figure 1.**
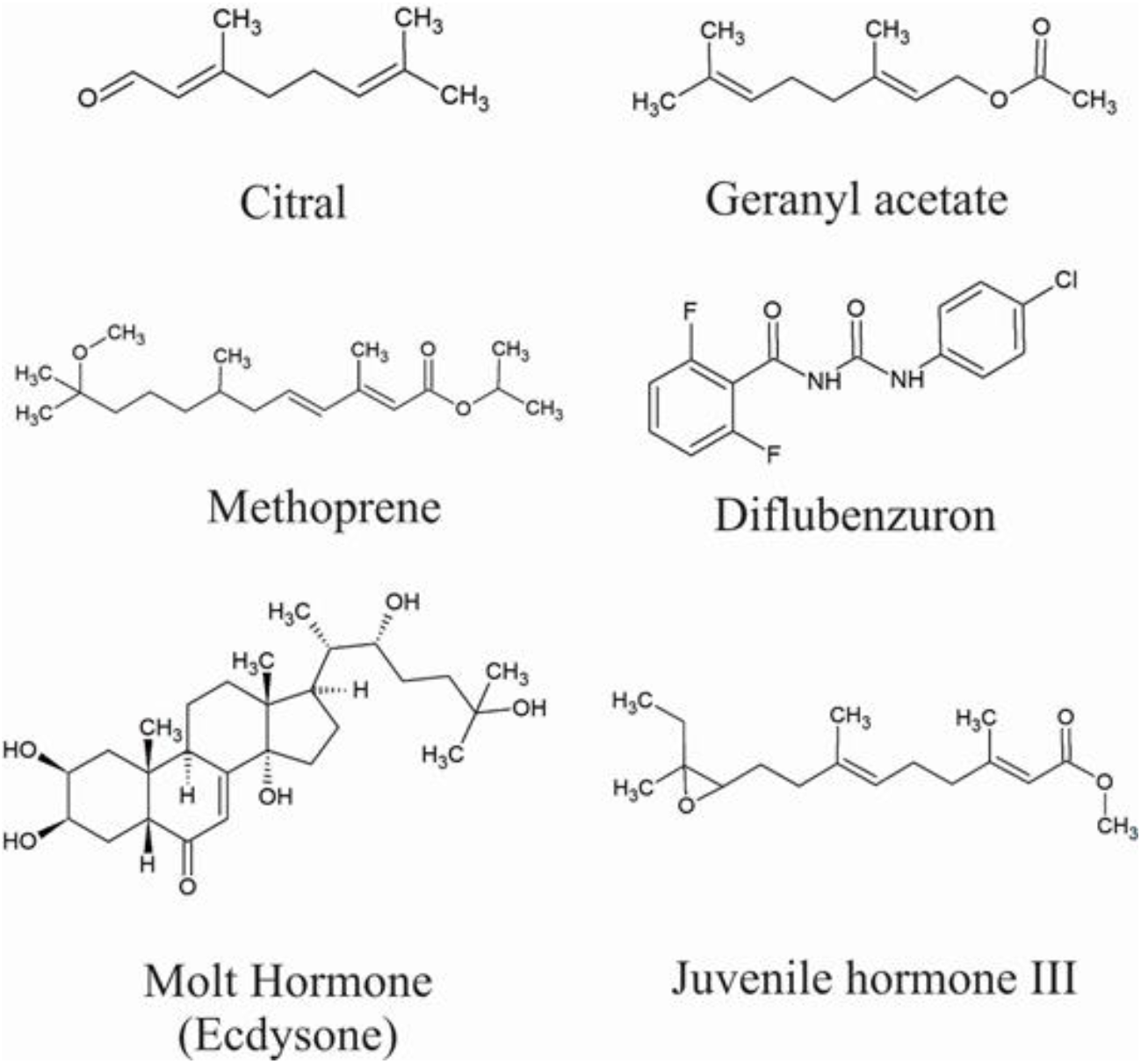
Chemical structures of the major compounds of *Cymbopogon flexuosus* EO (citral and geranyl acetate), the positive controls methoprene and diflubenzuron, and moult hormone (Ecdysone) and juvenile hormone III.

## Materials and methods

### Essential oil extraction and isolation of major compounds

*C. flexuosus* EO was provided and characterised by the ‘Centro de Investigación en biomoléculas (CIBIMOL) and Centro Nacional de Investigación para la Agroindustrialización de Plantas Aromáticas y Medicinales Tropicales (CENIVAM)’ of the Universidad Industrial de Santander (Colombia). The oil extraction methodology, as well as its chemical characterisation, was carried out following the methodology described by Stashenko *et al*. (2004). The essential oil was obtained by hydrodistillation (HD) and microwave-assisted hydrodistillation (MWHD). The components were identified by comparing their relative retention times and mass spectrometry with the standard compounds (Stashenko *et al*. 2004). The description of EO chemical characterisation was reported by Vera *et al*. (2014): citral (geranial 37.5% and neral 28.2%), geranyl acetate (10.0%), geraniol (9.0%), and β-Bourboneno trans-β-cariofilene (2.0%). The major compounds (citral and geranyl acetate) were commercially acquired from Sigma-Aldrich (St. Louis, MI).

### Biological material

The experiments were performed with a colony of *Ae. aegypti*, Rockefeller strain, maintained in an insectary at 25 ± 5 °C, with humidity of 70 ± 5% and photoperiod (12:12). Male adults were fed with a sugar solution of 10% honey according to the breeding model for this species (Pérez *et al*. 2004). Females were blood-fed with an albino Wistar rat *(Rattus norvegicus)* (WI IOPS AF/Han strain) provided by the bioterium of the Universidad Industrial de Santander. This research was approved by the Ethics Committee (CEINCI) (Minutes No. 22, Dec 6, 2019). Once the females were fed, glass cups with Whatman ^®^ # 1 filter papers folded as a cone were placed inside the breeding cages to internally coat the walls of the vessel and allow the collection of eggs. To synchronise egg hatching, the filter paper was removed after the embryo maturation process (72 hours), and then allowed to dry for three days at room temperature. Subsequently, hatching was stimulated by immersing the eggs in dechlorinated water. The larvae were kept in plastic trays and fed with TetraMin Tropical Flakes^®^ fish concentrate.

### Bioassays

#### Ovicidal activity

We used the methodology of Suman *et al*. (2013) to assess the ovicidal affectation. The sublethal concentration was established based on lethal concentrations (LC) calculated for *C. flexuosus* EO in the fourth early-stage larvae of *Ae. aegypti* by Vera *et al*. (2014) at 24 hours (CL_50_ = 17.2 mg.L^-1^; CL_95_ = 49.9 mg.L^-1^) and 48 hours (CL_50_ = 14.7 mg.L^-1^; CL_95_ = 55.6 mg.L^-1^), settling on 6, 18, and 30 mg.L^-1^.

We separated 20 gravid *Ae. aegypti* females in a security cage with dimensions of 40 x 40 x 40 cm. Inside the cage, we placed five plastic cups with 50 μL of EO dissolved in dimethyl sulfoxide (DMSO) at concentrations previously establish (6, 18, and 30 mg.L^-1^), a cup as a control treatment (49 mL of water + 1 mL of 0.5% DMSO), and a cup as a positive control (diflubenzuron at 6, 18, and 30 mg.L^-1^). To collect and count the eggs, each cup was coated inside with half of a Whatman ^®^ # 1 filter paper, folded in a cone shape. The oviposition lasted for eight consecutive days, changing the filter paper daily. Three replicates were used for each concentration, and the experiment was repeated on three different days. When the females oviposited, between 50 and 100 embryonated eggs (recollected after 72 hours) per cup were taken at random. The eggs were transferred to individual containers, according to the evaluated concentration, and they were counted and examined under a stereoscope to verify integrity. The hatching percentage of the eggs was evaluated up to 120 h after oviposition by submerging the eggs obtained in mineral water. The emerged first instar larvae were counted under a microscope, and eggs that did not hatch after seven days were considered not viable. The hatching value was estimated as the percentage of eggs that went on to the larval stage.

#### Larvicidal activity

For those experiments, we establish two sublethal concentrations of 3 and 6 mg.L^-1^ of *C. flexuosus* EO. To evaluate the major compounds citral and geranyl acetate, we also investigated the 3 and 6 mg.L^-1^ concentrations. However, we did not observe any effect on the larvae. For this reason, we increased the concentrations until establishing three sublethal concentrations of 18, 30, and 42 mg.L^-1^. Methoprene PESTANAL^®^ (Sigma-Aldrich) and diflubenzuron PESTANAL^®^ (Sigma-Aldrich) at 3 and 6 mg.L^-1^ were used as a comparison factor, and dimethylsulfoxide (DMSO 0.5%) as a negative control.

To determine the effect on larvae development, we used the methodology by Leyva *et al*. (2013) with some modifications. Ten larvae in stage L3 were selected and transferred by Pasteur pipettes to 200 mL plastic cups with 99.5 mL of chlorine-free water and 0.5 mL of each treatment (negative control, *C. flexuosus* EO, citral major component, geranyl acetate major component, and methoprene and diflubenzuron as positive controls).

All larvae treatments were supplied with 2% fish feed in chlorine-free water to ensure their survival. After 24 hours of treatment, the dead larvae were removed, and the survivors remained in the water until they pupated. At 48 hours, five larvae were taken for each treatment and were used to measure morphological parameters (in mm): larval length, cephalic diameter, and siphon length. All experiments were performed in triplicate and on three different days.

#### Effects on juvenile hormone and moulting hormone

To evaluate the effect of the juvenile hormone and molt hormone, 200 newly emerged larvae in stage L3 were selected and treated with *C. flexuosus* EO at the concentrations of 3 and 6 mg.L^-1^, and with citral and geranyl acetate at the concentrations of 18, 30, and 42 mg.L^-1^. For this experiment, molecular grade standards of methoprene PESTANAL^®^ (Sigma-Aldrich) and diflubenzuron PESTANAL^^®^^ (Sigma-Aldrich) were used (Figure 1).

After 24 hours, the larvae of each treatment were extracted and homogenised in 2 mL of a 50 mM phosphate buffer solution (pH 7.4) using a BeadBug TM Mini Homogeniser (Benchmark brand, model D1030). The homogenate obtained was filtered on glass wool to remove impurities and subsequently analysed by HPLC coupled with mass spectrometry. Six replicates of this experiment were performed with their respective calibration curve, using the molt hormone and juvenile hormone (Sigma-Aldrich) as a standard. We adapted the methodology of Zhou *et al*. (2011) for the quantification of the juvenile hormone (JH) and moult hormone (HM) by Liquid Chromatography coupled with mass spectrometry (ESI - IT). An Elite LabChrom liquid chromatography (Hitachi-VWR) with a detection range of 190-600 nm (UV), injection volume 0.1-90 μL, and flow range 0.001-10 mL/min coupled to a mass spectrometer (Electrospray ionisation - Ion trap analyser (ESI - IT)), with an Octadecylsiloxane-ODS column Zorbax XDB-C18 (4.6 mm x 10 mm, 2.0μm) was used.

Water (0.1% formic acid) plus acetonitrile (0.1% formic acid) and water (0.1% formic acid) plus methanol (0.1% formic acid) were used as the mobile phase, applying a binary gradient at a flow of 0.30 mL/min, with an injection volume of 10 μL. The total running time of the equipment was 15 min. The retention time corresponding to the signal of each hormone was obtained, and the concentration levels of the hormones in each treatment were determined.

#### Statistical analyses

All data were subjected to normality tests. When the data presented a normal distribution, we used an ANOVA and subsequently, Tukey’s test. If the distribution was not normal, non-parametric tests were applied and subsequently the Kruskal-Wallis test. Statistical significance with values of *p* ≤ 0.05. The results were analysed with Statistica Software V11.

## Results

### Eggs activity

We obtained a 65% decrease in hatching of eggs treated with *C. flexuosus* EO at concentrations of 6, 18, and 30 mg.L^-1^. The minor hatching percentage was at 23.3% at 18 mg.L^-1^, followed by 34.6% at 30 mg.L^-1^, and 45.3% at 6 mg.L^-1^. Significant differences were found when comparing the hatching percentages at different concentrations of *C. flexuosus* EO and the control group: at 18 mg.L^-1^ (ANOVA: H (1, N = 12), *p* = 0.017 and 30 mg.L^-1^ (ANOVA: H (1, N = 12), *p* = 0.038) (Figure 2). The eggs treated with diflubenzuron had hatching percentages at upper 80% in all concentrations used (6 mg.L^-1^ = 85.6%, 18 mg.L^-1^ = 85.6%, and 30 mg.L^-1^ = 84.4%) (Figure 2). No significant differences were found between diflubenzuron concentrations and the control group hatching percentages (Kruskal-Wallis test: H (3, N = 12) = 5.290476, *p* = 0.1517). Significant differences were found, in all concentrations, when comparing the hatching percentages of *C. flexuosus* EO and diflubenzuron (ANOVA: H (1, N = 6), *p* < 0.05) (Figure 2).

**Figure 2.**
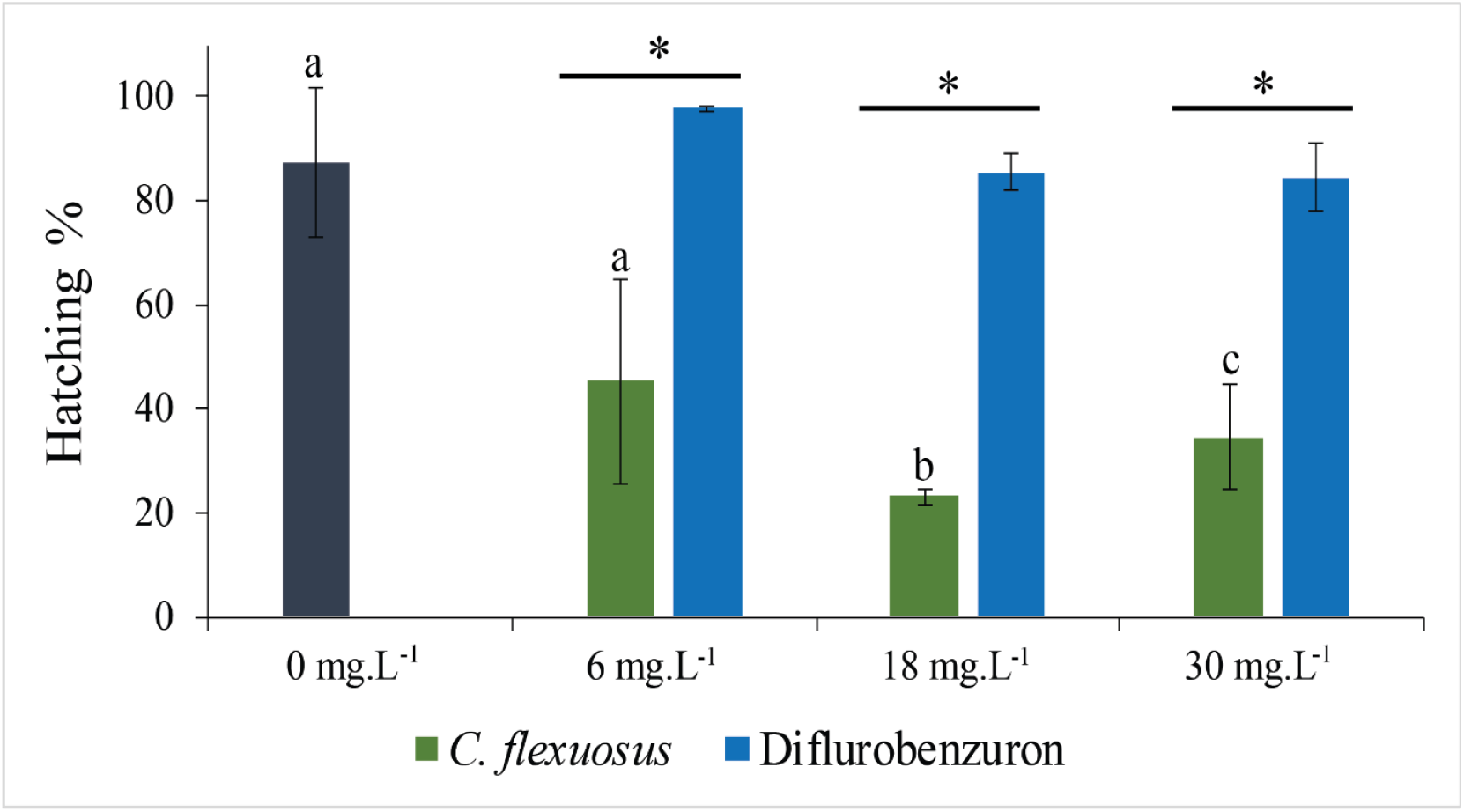
Hatching percentages of *Ae. aegypti* eggs subjected to different concentrations of *C. flexuosus* EO and diflubenzuron (6 mg. L^-1^, 18 mg. L^-1^, and 30 mg. L^-1^). Different letters (a, b, and c) indicate significant statistical differences between 18 mg.L^-1^ and 30 mg.L^-1^ EO concentrations and the control group (Tukey’s test *p* ≤ 0.05). (*) Significant statistical differences between each concentration of *C. flexuosus* and diflubenzuron (Tukey’s test *p* < 0.05).

### Larvicidal activity

The larvae treatments with *C. flexuosus* EO, diflubenzuron, and methoprene showed changes in larvae length, cephalic diameter, and siphon length (Figure 3). We observed a significative (ANOVA: H (2, N = 9), *p* ≤ 0.05) increase in the head width in larvae treated with *C. flexuosus* EO at concentrations of 3 mg.L^-1^ (0.8 ± 0 mm) and 6 mg.L^-1^ (0.8 ± 0.0 mm) compared with the head width average in the control group (0.7 ± 0.0 mm). The length of larvae increases significantly (ANOVA: H (2, N = 9), *p* ≤ 0.05) with *C. flexuosus* EO at concentrations of 3 mg.L^-1^ (6 ± 0.0 mm) and 6 mg.L^-1^ (6 ± 0.7 mm) compared with the head width average in the control group (5.3 ± 0 mm). No changes were observed in the siphon length subjected to different concentrations of *C. flexuosus* EO (3 mg.L^-1^ = 0.7 ± 0 mm; 6 mg.L^-1^ = 0.7 ± 0 mm) compared with the head width average in the control group (0.7 ± 0 mm) (ANOVA: H (2, N = 9), *p* ≥ 0.05) (Figure 4).

**Figure 3.**
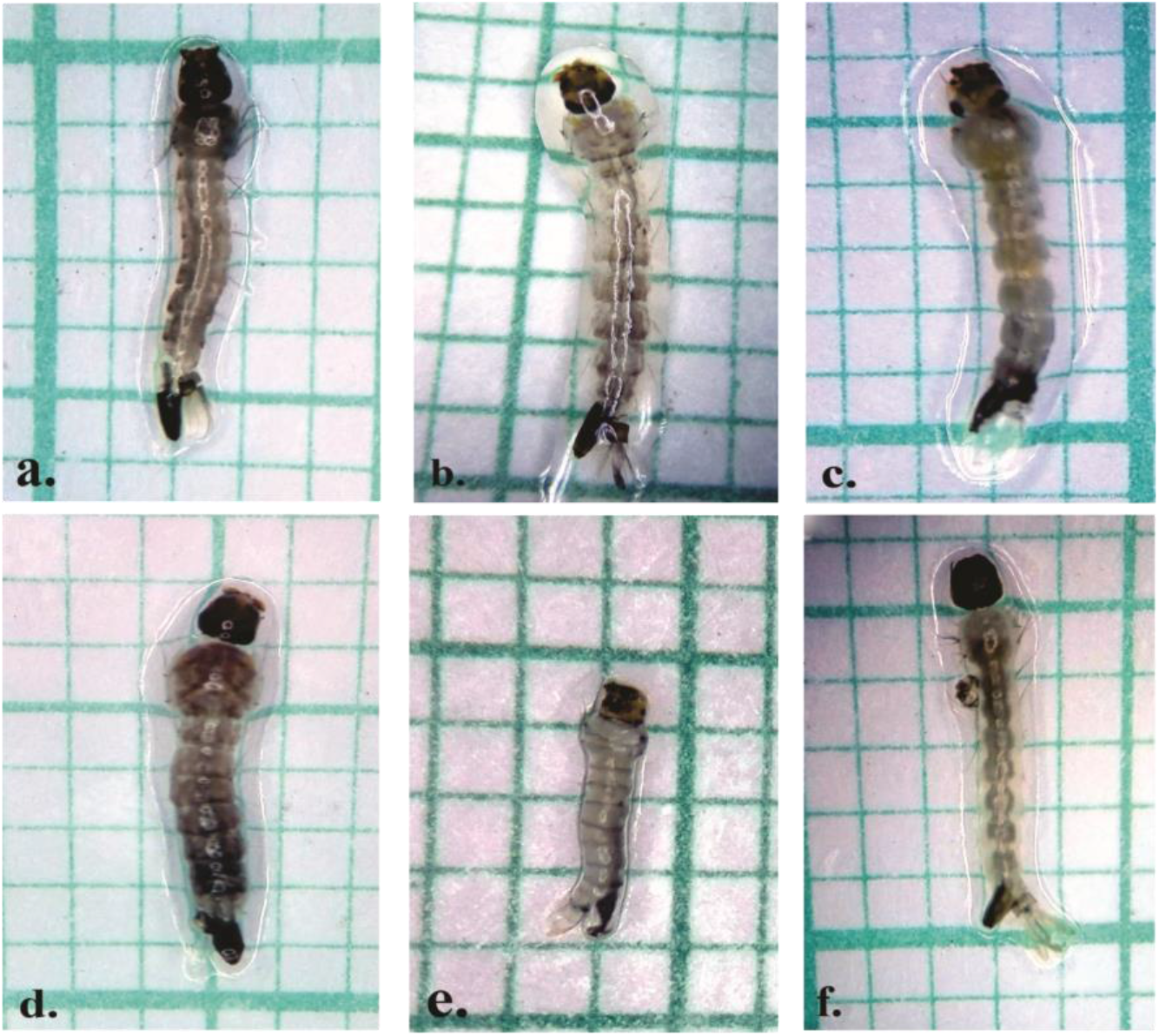
Morphological differences in *Ae. aegypti* larvae under different treatments. a. Control group; b. *C. flexuosus* EO; c. citral; d. geranyl acetate; e. diflubenzuron; and f. methoprene.

**Figure 4.**
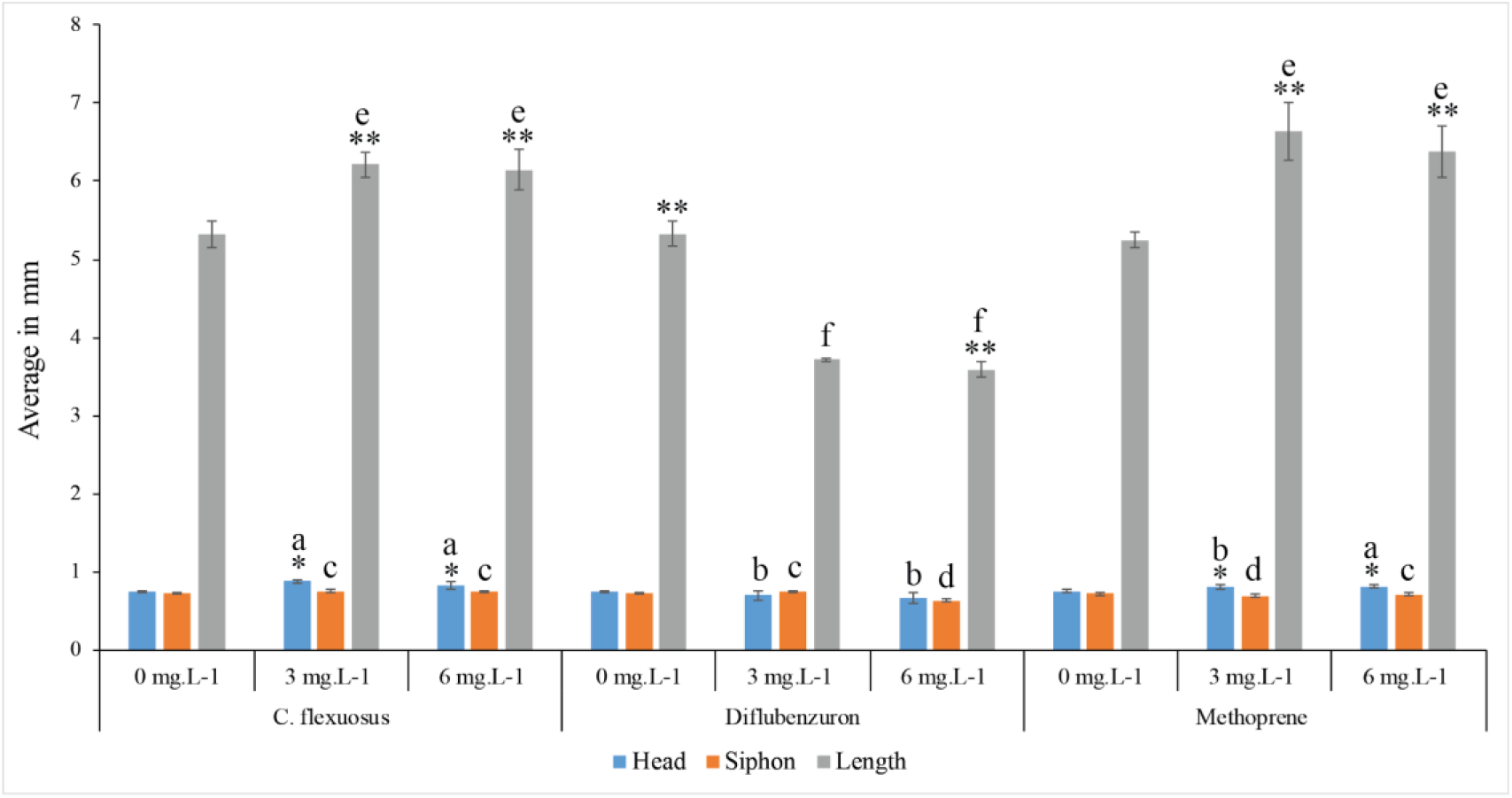
Effect of different concentrations of *C. flexuosus* EO (3 mg.L^-1^ and 6 mg.L^-1^), Diflubenzuron and methoprene in the development of *Ae. aegypti* larvae. Measures of larvae length, cephalic diameter, and siphon length presented in average millimetres. Different letters indicate significant statistical differences between (a, b) each treatment by cephalic diameter without a control group (Tukey’s test *p* ≤ 0.05), (c, d) each treatment by siphon length without a control group (Tukey’s test *p* ≤ 0.05), (e, f) each treatment by length larvae without control group (KW test p ≤ 0.05; Tukey’s test *p* ≤ 0.05). (*) Significant differences by each treatment between concentrations and the control group (Tukey’s test *p* ≤ 0.05) by cephalic diameter. (**) Significant differences by each treatment between concentrations and the control group (Tukey’s test *p* ≤ 0.05) by length larvae.

*Ae. aegypti* larvae treated with diflubenzuron showed a decrease in the morphological parameters measured. We observed a significant decrease in the head width of larvae treated with diflubenzuron at concentrations of 3mg.L^-1^ (0.7 ± 0.1 mm) and 6 mg.L^-1^ (0.7 ± 0.1 mm) compared with the control group (0.7 ± 0.0 mm) (KW test: H (2, N = 9) =2.509804, *p* = 0.2851). Also, we noted a small decrease in siphon length under the diflubenzuron concentrations used (3mg.L^-1^ = 0.7 ± 0.0 mm and 6 mg.L^-1^ = 0.6 ± 0.0 mm) compared with the control group (0.7 ± 0.0 mm), without significant differences with the control group (KW test: H (2, N = 9) = 5.600000, *p* = 0.0608). Larvae length showed a decrease with diflubenzuron concentrations of 3 mg.L^-1^ (3.7 ± 0.0 mm) and 6 mg.L^-1^ (3.6 ± 0.1 mm) compared with the control group (5.3 ± 0.2 mm), with significant differences between a 6 mg.L^-1^ concentration and the control group (KW test: H (2, N=9) = 6.937853, *p* = 0.0312) (Figure 4).

The morphological parameters of larvae treated with methoprene showed a small increase when compared with the control group. The measured head width increased at concentrations of 3 mg.L^-1^ and 6 mg.L^-1^ (0.8 ± 0.0 mm and 0.8 ± 0.0 mm, respectively) compared with the control group (0.7 ± 0.0 mm) (ANOVA: H (2, N=9), *p* ≤ 0.05) (Figure 3). Larvae length significantly increased with methoprene concentrations of 3 mg.L^-1^ (6.6 ± 0.4 mm) and 6 mg.L^-1^ (6.4 ± 0.3 mm) compared with the control group (5.2 ± 0.1 mm) (ANOVA: H (2, N = 9),*p ≤* 0.05). The measures of siphon length after methoprene treatment at 3 mg.L^-1^ (0.7 ± 0.0 mm) and 6 mg.L^-1^ (0.7 ± 0.0 mm) were similar to the control group (0.7 ± 0.0 mm), without significant differences (ANOVA: H (2, N = 9), *p* ≥ 0.05) (Figure 4). We compared each morphological measure by the substance tested and its concentration. Significant differences were found for head width measured at a concentration of 3 mg.L^-1^ among *C. flexuosus* EO with diflubenzuron and methoprene (ANOVA: H (2, N = 9), *p* ≤ 0.05). At a concentration of 6 mg.L^-1^, we found significant differences between *C. flexuosus* EO and diflubenzuron (ANOVA: H (2, N = 9), *p* ≤ 0.05) (Figure 4).

For siphon length, statistical differences were found at a concentration of 3 mg.L^-1^ between *C. flexuosus* EO - Diflubenzuron (ANOVA: H (2, N = 9), *p* ≤ 0.05), and between Diflubenzuron - Methoprene (ANOVA: H (2, N = 9),*p ≤* 0.05). At a concentration of 6 mg.L^-1^, statistical differences were found between *C. flexuosus* EO - Diflubenzuron (ANOVA: H (2, N = 9),*p* ≤ 0.05), and between Diflubenzuron - Methoprene (ANOVA: H (2, N = 9), *p* ≤ 0.05) (Figure 4). For larvae length, statistical differences were found at a concentration of 3 mg.L^-1^ between Diflubenzuron and Methoprene (KW test: H (2, N = 9) = 6.543417 *p* = 0.0379). At a concentration of 6 mg.L^-1^ statistical differences were found between *C. flexuosus* EO - Diflubenzuron (ANOVA: H (2, N = 9),*p* ≤ 0.05), and between Diflubenzuron - Methoprene (ANOVA: H (2, N = 9), *p* ≤ 0.05) (Figure 4).

Given the average increase of the morphological measures of *Ae. aegypti* larvae treated with *C. flexuosus* EO, we evaluated the effect of their major compounds (citral and geranyl acetate) at concentrations of 18, 30, and 42 mg. L^-1^. We did not observe a significant decrease in the cephalic diameter with citral (30 mg.L^-1^ = 0.7 ± 0.0 mm; 42 mg.L^-1^ = 0.7 ± 0.0 mm) (ANOVA: H (3, N = 12), *p* ≤ 0.05) and geranyl acetate (30 mg.L^-1^ = 0.7 ± 0.0 mm; 42 mg.L^-1^= 0.7 ± 0.0 mm) (KW test: H (3, N = 12) = 7.228944, *p* = 0.0649) compared to the control group (0.8 ± 0.0 mm) (Figure 5).

**Figure 5.**
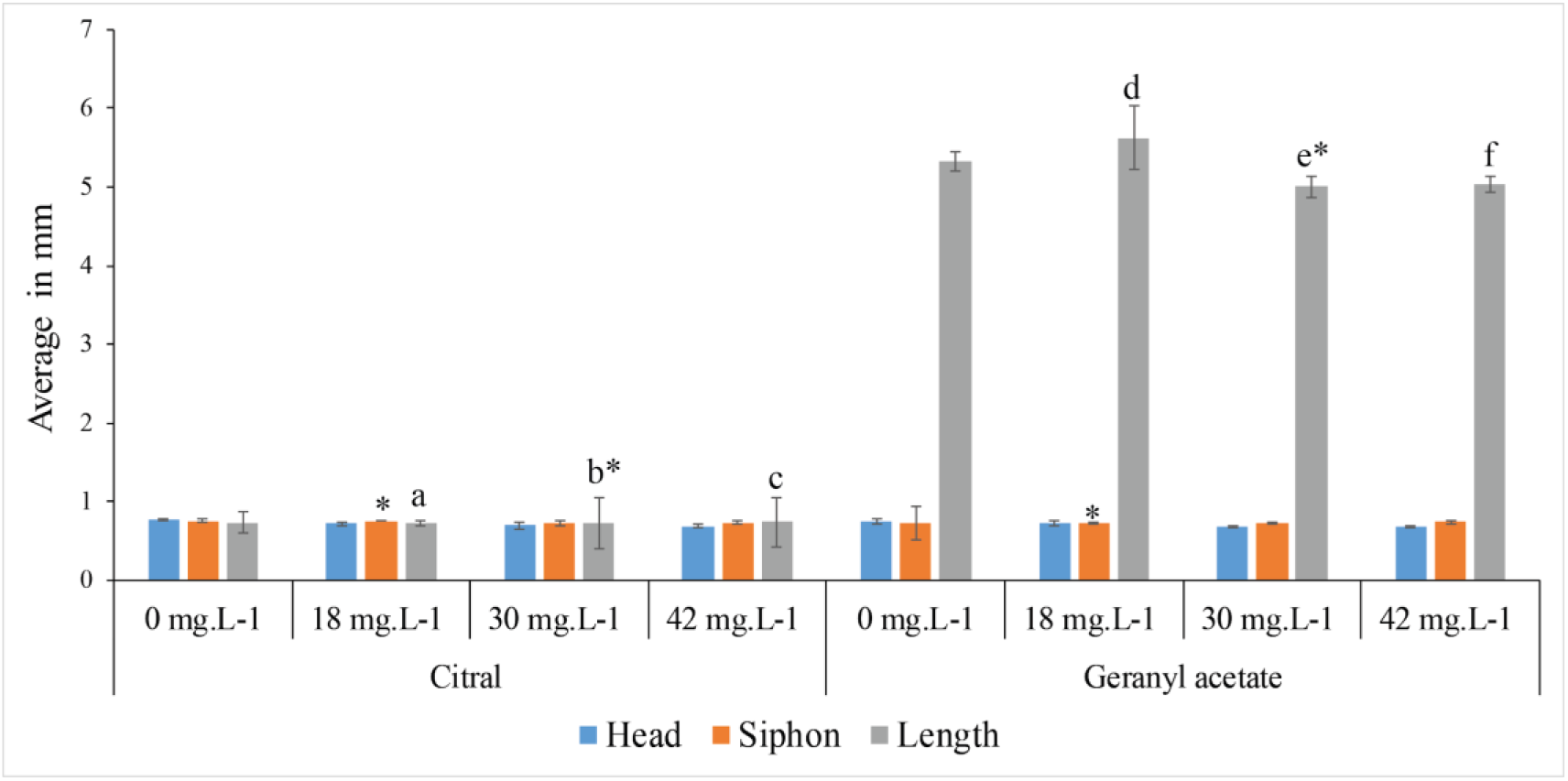
Effect of major compounds of *C. flexuosus* EO (citral and geranyl acetate) at a concentration of 18, 30, and 42 mg.L^-1^ over the larvae development of *Ae. aegypti*. Measures of larvae length, cephalic diameter, and siphon length present in average millimetres. Different letters (a, b, c, d, e, f) indicate significant differences among concentrations for each substance and the control group (Tukey’s test *p* ≤ 0.05). (*) Significant differences between citral and geranyl acetate by siphon length (Tukey’s test *p* ≤ 0.05); between citral and geranyl acetate by length larvae (Tukey’s test *p* ≤ 0.05).

For siphon length, we did not observe a significant decrease with the major compound citral at concentrations of 30 mg.L^-1^ (0.7 ± 0.0 mm) and 42 mg.L^-1^ (0.7 ± 0.0 mm) compared to the control group (0.7 ± 0.0 mm) (KW test: H (3, N=12) = 4.714286, *p* = 0.1940) (Figure 3). With the major compound geranyl acetate, we did not obtain significant differences for siphon length at a concentration of 42 mg.L^-1^ (0.7 ± 0.0 mm) compared to the control group (0.7 ± 0.2 mm) (ANOVA: H (3, N = 12), *p* ≤ 0.05) (Figure 5).

For total larvae length, we observed an increase with the major compound citral at concentrations of 18 mg.L^-1^ (5.6 ± 0.0 mm) and 30 mg.L^-1^ (5.8 ± 0.3 mm) compared to the control group (5.4 ± 0.1 mm). Additionally, we found significant differences among the concentration of 42 mg.L^-1^ with 18 mg.L^-1^ and 30 mg.L^-1^ (ANOVA: H (3, N = 12),*p* ≤ 0.05). With the major compound geranyl acetate, we observed a significant increase (ANOVA: H (3, N = 12), *p ≤* 0.05) at the concentration of 18 mg.L^-1^ (5.6 ± 0.0 mm), and a decrease at concentrations of 30 mg.L^-1^ (5 ± 0.1 mm) and 42 mg.L^-1^ (5 ± 0.1 mm) in the total larvae length, when compared to the control group (5.4 ± 0.1 mm) (Figure 5).

### Inhibitory activity of development

*C. flexuosus* EO (3 and 6 mg. L^-1^) exhibits an inhibitory effect on the *Ae. aegypti* development. None of the L3 larvae reached the pupa stage; they remained in the larval phase for 12 days (Figure 6). This performance was similar to the results obtained with positive controls (diflubenzuron and methoprene) (Figure 7). On the other hand, 60% of the larvae in the control group reached the pupa stage at the seventh day of development, and the life cycle was completed on the twelve-day (Figures 6 and 7).

**Figure 6.**
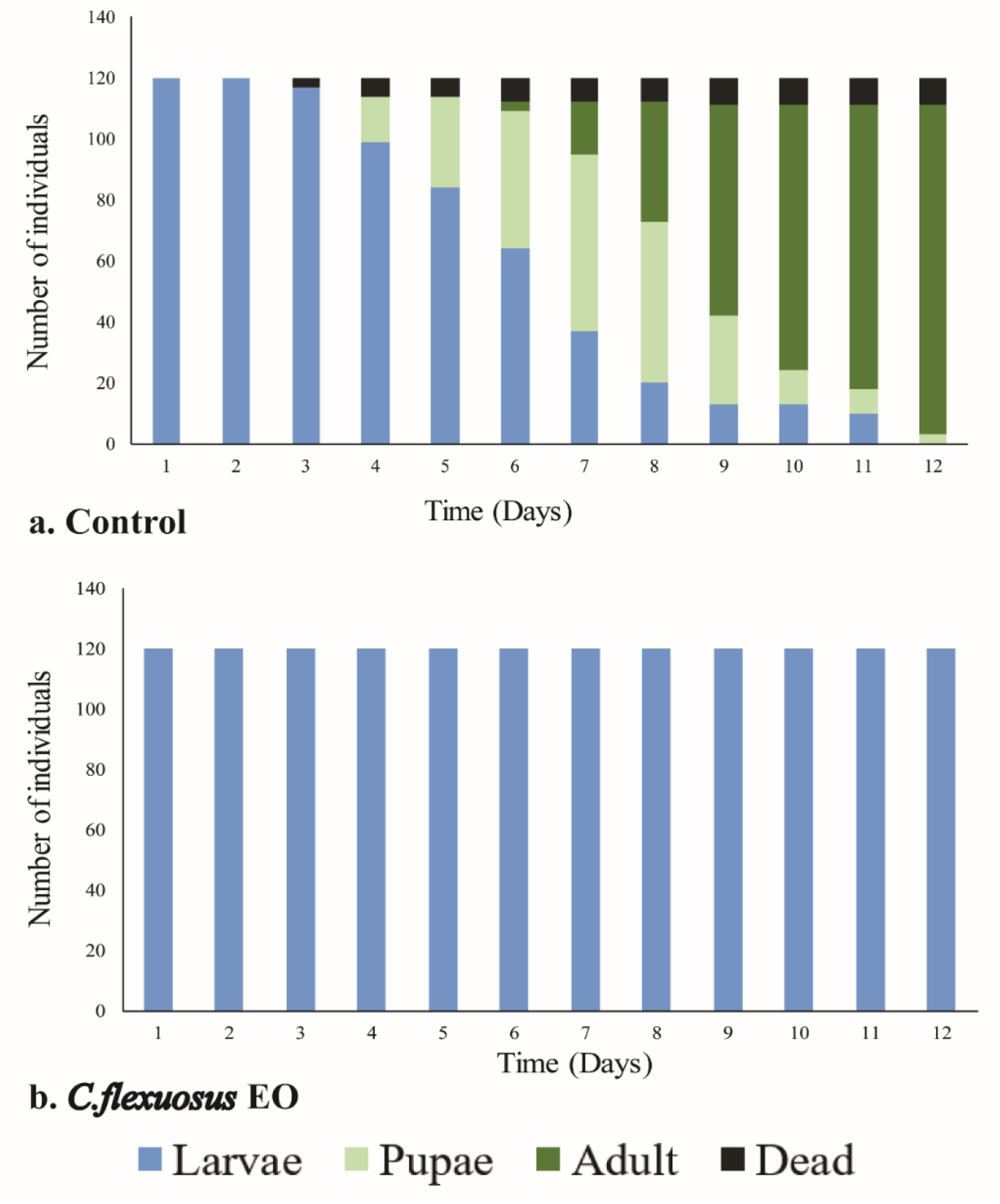
Duration in days of larval development (L3 and L4), pupae, and adult stages of *Aedes aegypti*. The individuals were treated with *Cymbopogon flexuosus* EO at a concentration of 3 mg.L^-1^ and 6 mg.L^-1^.

**Figure 7.**
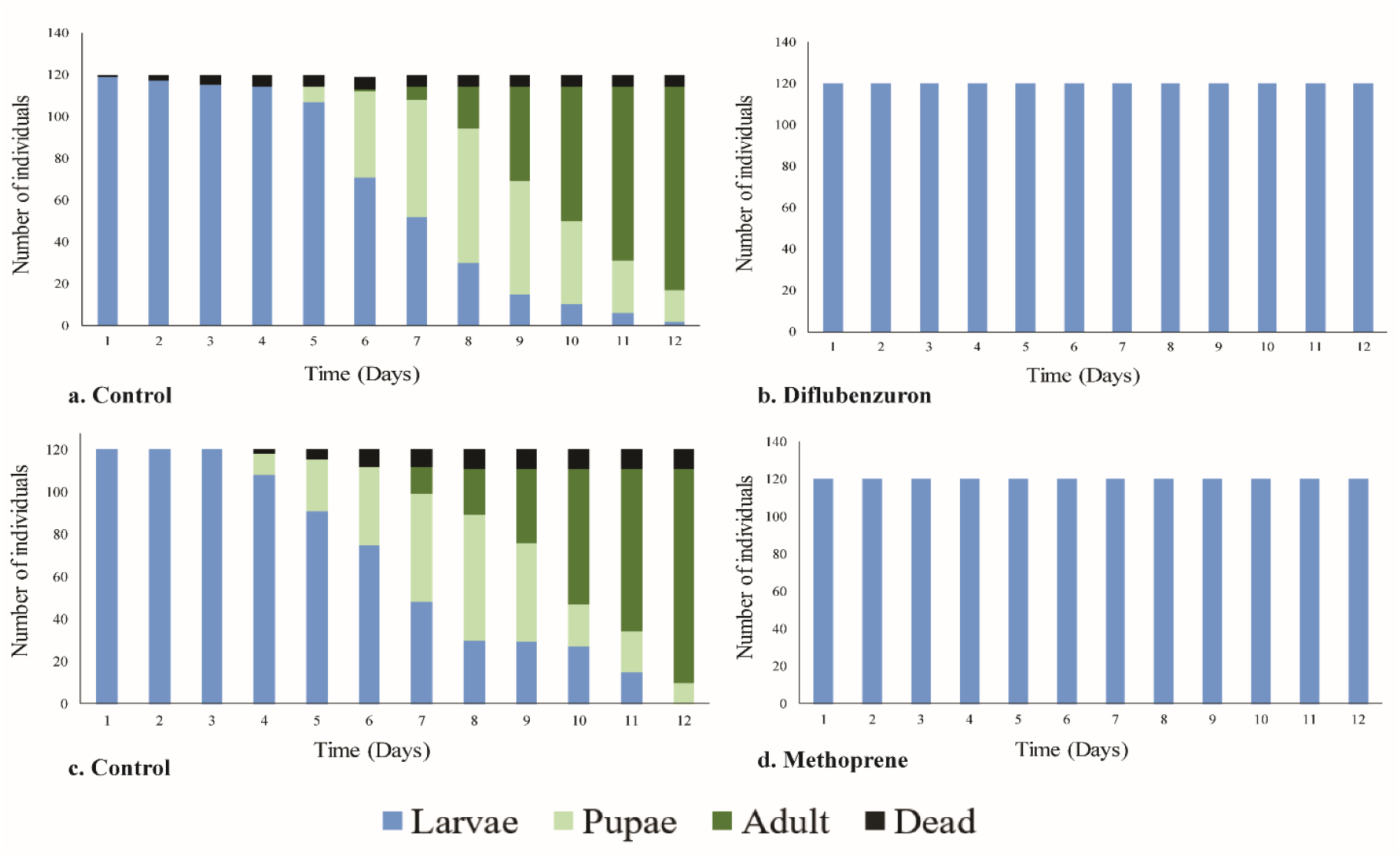
Duration in days of larval development (L3 and L4), pupae, and adult stages of *Aedes aegypti*. The individuals were treated with growth regulators at concentrations of 3mg.L^-1^ and 6 mg.L^-1^. **a-c.** Control treatments. **b.** Diflubenzuron treatment. **c.** Methoprene treatment.

The major compound citral (18 and 30 mg. L^-1^) exhibited similar results to the control group. The life cycle was completed on the twelve-day, and 60% of the larvae reached the pupa stage at the seventh day (Figure 8). However, we obtained 100% mortality at a concentration of 42 mg.L^-1^ on the fifth-day of exposure without reaching the pupa stage (Figure 8). The major compound geranyl acetate exhibited similar results in the time of life cycle with the control group at concentrations of 18 and 30 mg.L^-1^. The individuals reached the adult stage on the twelfth day, and 65% of the larvae reached the pupa stage (Figure 9). With the major concentration (42 mg.L^-1^), we obtained 100% mortality in the larvae at the first day of treatment (Figure 7).

**Figure 8.**
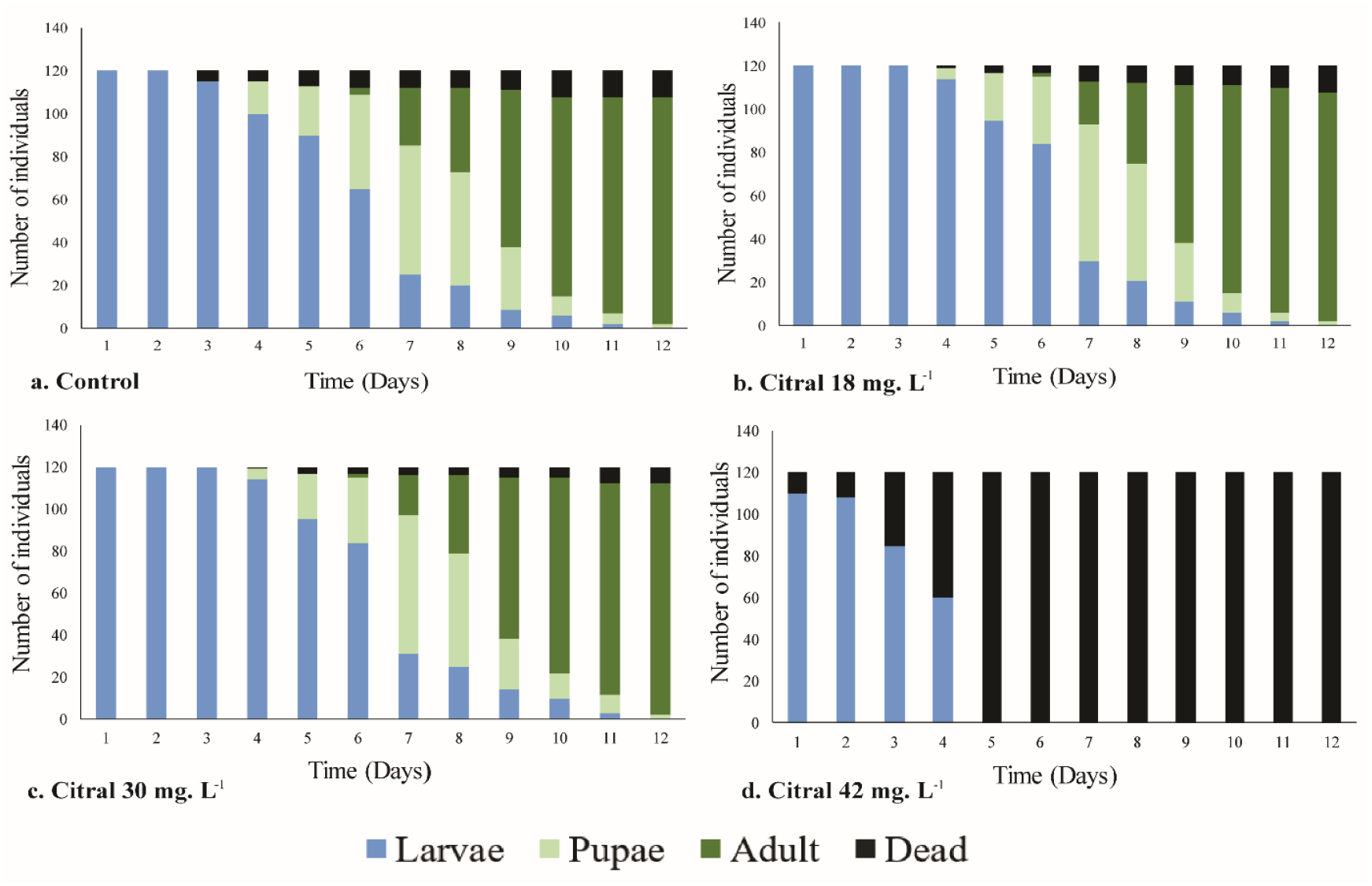
Duration in days of larval development (L3 and L4), pupae, and adult stages of *Aedes aegypti*. The individuals were treated with the major compound citral. **a.** Control treatments. **b.** Treatment at a concentration of 18 mg.L^-1^. **c.** Treatment at a concentration of 30 mg.L^-1^. **d.** Treatment at a concentration of 42 mg.L^-1^.

**Figure 9.**
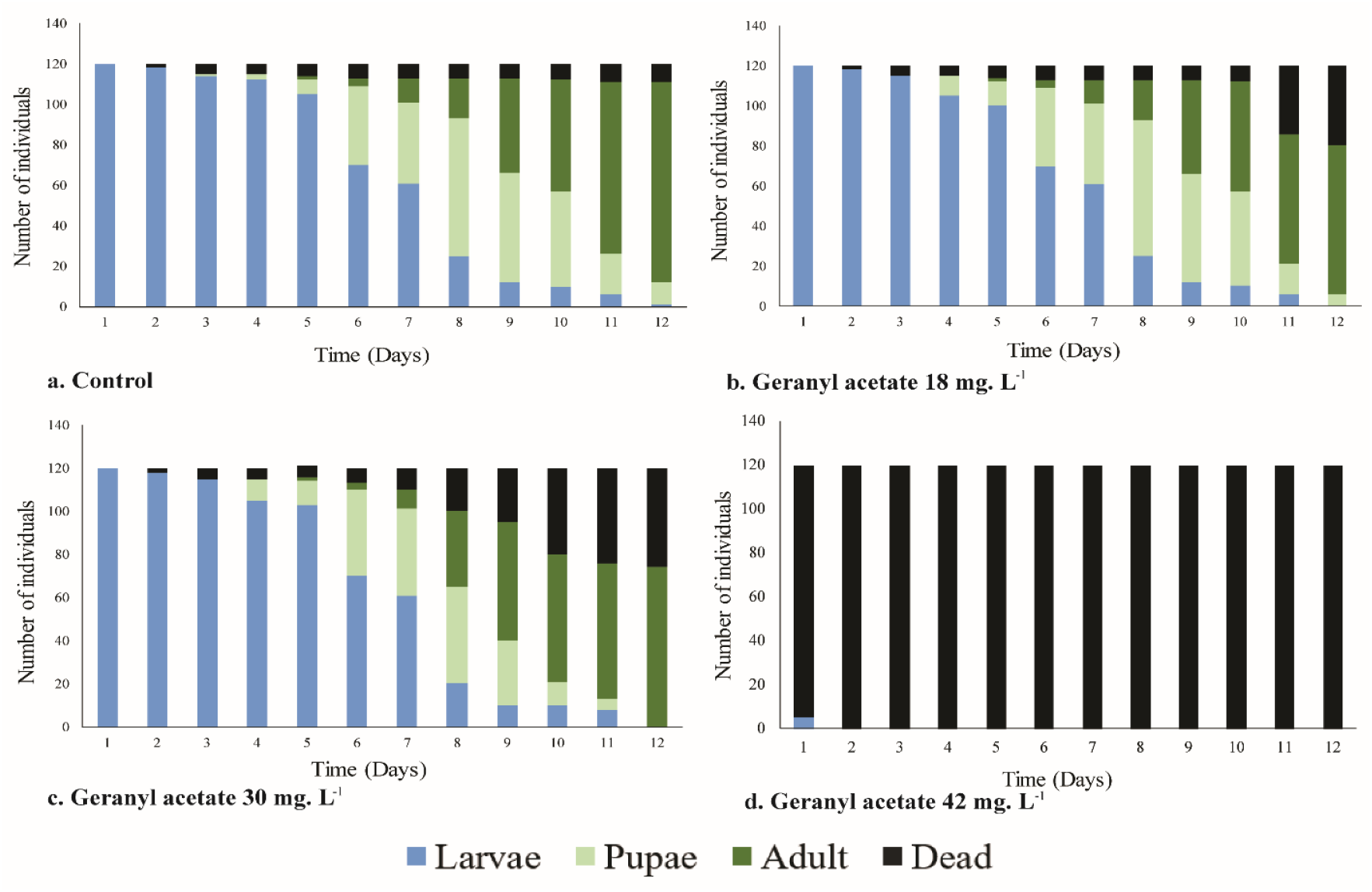
Duration in days of larval development (L3 and L4), pupae, and adult stages of *Aedes aegypti*. The individuals were treated with the major compound geranyl acetate. **a.** Control treatments. **b.** Treatment at a concentration of 18 mg.L^-1^**. c.** Treatment at a concentration of 30 mg.L^-1^. **d.** Treatment at a concentration of 42 mg.L^-1^.

### Juvenile hormone and moult hormone levels in larvae treated with EO

We measured juvenile hormone III (JH III) and moult hormone (MH) from larvae (L4) by HPLC coupled with mass spectrometry using methanol as a mobile phase. The MH was followed at 6.77 min retention time (Supplementary Fig. 1a) that corresponds to the molecular ion (mass/H^+^ of 481.1 g.mol^-1^). The JH III was followed at 9.86 min retention time (Supplementary Fig. 2a) that corresponds to 266.6 g.mol^-1^ from molecular ion mass.

Also, we made chromatograms of *Ae. aegypti* larvae (L4) treated with positive controls methoprene and diflubenzuron and compared the peaks found by MH and JH III. With methoprene, we observed greater value of areas under the curve for the MH (82 ± 3.4% at 3 mg.L^-1^ and 65 ± 0% at 6 mg.L^-1^) in contrast with JH III (18 ± 3.4% at 3 mg.L^-1^ and 35 ± 0% at 6 mg.L^-1^). We observed similar values with diflubenzuron for the MH (80 ± 0% at 3 mg.L^-1^ and 81 ± 0% at 6 mg.L^-1^) and JH III (20 ± 0% at 3 mg.L^-1^ and 19 ± 0% at 6 mg.L^-1^) (Supplementary Fig. 3).

To analyse the chromatograms of *Ae. aegypti* larvae (L4) treated with *C. flexuosus* EO and its major compounds (citral and geranyl acetate), we compared the values of areas under the curve observed for MH and JH III in the positive controls (methoprene and diflubenzuron) with the values observed with *C. flexuosus* EO (MH: 30 ± 1.1% at 3 mg.L^-1^ and 43 ± 0% at 6 mg.L^-1^; JH III: 4 ± 2.5% at 3 mg.L^-1^ and 22 ± 0% at 6 mg.L^-1^), citral (MH: 26 ± 0% at 42 mg.L^-1^; JH III: 1 ± 0% at 42 mg.L^-1^), and geranyl acetate (MH: 36 ± 0% at 30 mg.L^-1^ and 29 ± 0% at 18 mg.L^-1^; JH III: 3 ± 0% at 30 mg.L^-1^ and 6 ± 0% at 18 mg.L^-1^). The minor values of areas under the curve obtained with *C. flexuosus* EO and its major compounds (citral and geranyl acetate) treatment indicate the effects on the concentration of MH and JH III in the larvae of *Ae. aegypti*, with values under the detection limit for this technique (Supplementary Fig. 4).

## Discussion

The action mechanism of most synthetic pesticides explains the interaction or union of a synthetic molecule with a specific target from a protein or biomolecule. This union or interaction triggers a series of biochemical events that modify the physiological functions or cause death (Casida 2009). Although the synthetic pesticide has four or six targets, the majority act directly on neuronal receptors or ions channels. The action on these targets makes them highly specific, contributing to the generation of resistance mechanisms (Narahashi *et al*. 2007, Casida 2009).

The search of new molecules that counter the resistance levels has prompted the screening of substances of natural origin like essential oils and major compounds (Casida 2009, Liu 2015). The importance of these natural substances resides in a wide range of molecular targets on proteins (enzymes, receptors, ions channels, structural proteins), nucleic acids, biomembranes, and interfered on different metabolic pathways (Rattan 2010).

### Eggs activity

One of the major problems in the *Ae. aegypti* control resides in that its eggs are highly resistant to desiccation periods, remaining in latency near to the end of embryonic development (Rezende *et al*. 2008). There are scarce studies that mentioned the ovicidal actions of natural substances. Plants of the genus *Cymbopogon* have been recognised by their ovicidal effect against different species of mosquitoes. Warikoo *et al*. (2011) reported 100% of hatching inhibition with *C. nardus* EO on *Ae. aegypti* eggs to concentrations of 1%, 10%, and 100 %. Additionally, Pushpanathan *et al*. (2006) obtained 22.4% of mortality in *Culex quinquefasciatus* eggs treated with 100 mg L^-1^ of *C. citratus*, and the mortality of eggs increased to 100% with a concentration of 300 mg.L^-1^. The results obtained by these authors are consistent with the results observed in the present study, where hatching decreased in *Ae. aegypti* eggs treated with sublethal concentrations (6 mg. L^-1^, 18 mg. L^-1^, and 30 mg.L^-1^) of *C. flexuosus* EO.

*Ae. aegypti* eggs are characterised by two layers, the external (exo-chorion) and internal (endochorion). On the inner side of the endo-chorion is a serosal cuticle, which is formed during the embryogenesis process and protects the embryo from external factors like desiccation or the presence of bacteria or insecticides (Rezende *et al*. 2008). However, this serosal cuticle can be disrupted by lipophilic substances (Rajkumar *et al*. 2011) like terpenes citral and geranyl acetate, major compounds of *C. flexuosus* EO. The hatching inhibition by these terpenes is due to an interruption of embryo development caused by physiological alterations in water and gas exchanges, enzymatic modifications, and hormonal changes. These physiological alterations cause the embryo to not fully develop, decreasing the percentage of hatching (Saxena and Sharma 1972, Rajkumar *et al*. 2011), as observed in the results of the present study (Figure 2).

Additionally, our hatching inhibition values obtained with diflubenzuron (close to 20%) are consistent with those reported by Suman *et al*. (2013), who treated freshly laid and embryonated *Ae. aegypti* eggs with concentrations of 0.001, 0.01, 0.1, and 1 mg. L^-1^, and showed a hatching inhibition of 25.5%. Although diflubenzuron is a chitin synthesis inhibitor in the larval and pupal stages, the hatching inhibition values observed with this substance are less than the values obtained with *C. flexuosus* EO (Figure 2). The mechanism of action of this insecticide is related to the inhibition of transmembrane transport of chitin precursors, preventing the formation of microfibrils and embryo development (Reynolds 1987).

### Activity on larvae and the life cycle

We found that the sub-lethal concentrations of *C. flexuosus* EO affect the normal development of the juvenile phases of *Ae. aegypti*. The EO caused development alterations and morphological changes in larvae, represented in the lack of change in stage, an increase in the size of the larva, the width of the head, and length of the siphon (Figure 3 and 4). These results agree with the report by Soonwera and Phasomkusolsil (2016), who found morphological abnormalities (in *Ae. aegypti* larvae, pupae, and adults), and high percentages of larvae mortality that do not reach the pupae stage under concentrations of 1, 5, and 10% of *C. citratus* EO.

We observed similar morphological changes in larvae treated with *C. flexuosus* EO and methoprene, with an increase in the width of the head and the size of the larva (Figure 3 and 4). Because methoprene is a juvenile hormone analogue, it acts directly on the moult, permitting the increase of larvae size but preventing the change to pupae stage (Castro *et al*. 2007). This inhibition effect on the juvenile stage change was also observed in larvae treatment with *C. flexuosus* EO, with a maximum duration period in the larval stage of 12 days. On the other hand, the measures of the morphological parameters with diflubenzuron treatment were lower than measures in the presence of *C. flexuosus* EO (Figure 6). Diflubenzuron inhibits chitin synthesis, causing alterations in the cuticle layers during the moult (Viñuela *et al*. 1991, Hoffmann and Lorenz 1998). These alterations also alter the normal duration of development stages. In our case, they caused larval development inhibition, which was a duration greater than 12 days of observation.

To verify if the effect observed by *C. flexuosus* EO is due to the action of the major compounds (citral and geranyl acetate), we found that the increase of the size of the larva was the unique parameter with a similar change to the effect caused by EO. Related to larval stage duration, we did not observe a notable inhibition at concentrations of 18 and 30 mg.L^-1^. However, at the higher concentration (42 mg.L^-1^), both compounds caused larval mortality (Figure 8 and 9).

The major compound in *C. flexuosus* EO is citral. It is a monoterpene with an acyclic aldehyde functional group, responsible from the aroma in species from the genus *Cymbopogon*, with different biological properties. The mortality per cent obtained in the present study with citral is similar as reported by other studies, with a mortality per cent of 100% in *Ae. aegypti* larvae (Figure 8) (Freitas *et al*. 2010, Ganjewala *et al*. 2012). Concerning size modifications in larvae, the mechanism of action involved may be related to the study by Chaimovitsh *et al*. (2010). These authors reported that the principal target of citral is the microtubules, which directly interact with tubulin and cause damage in the cell membrane in both animal and vegetable cells. Additionally, Orhan *et al*. (2008) found that citral is a reversible competitive inhibitor of acetylcholinesterase (AChE), affecting nerve impulse transmission. Finally, Matsuura *et al*. (2006) found that citral has an inhibitory activity against tyrosinase, an enzyme responsible for different biological process, among those found the exoskeleton consolidation in arthropods during the molt process (Tareq *et al*. 2006).

Concerning the bioactivity observed under geranyl acetate treatment (an acyclic monoterpene), the results of the present study are similar to those reported by Michaelakis *et al*. (2014) who evaluated the repellent and larvicide activity on *Ae. aegypti* of different compounds and derivatives. They argue that although the larvicidal activity of geranyl acetate is minor to that of citral, geranyl acetate presented higher repellent and larvicide activity related to his functionality and degree of saturation. Additionally, Cheng *et al*. (2009) reported a larvicide activity of 100% with geranyl acetate against *Ae. albopictus* larvae. Although the exact mechanism of action involved is unknown, many authors mentioned that the monoterpenes structure and their functional modifications play a fundamental role in the activity against mosquitoes, potentialised in acetylated forms (Pandey *et al*. 2013).

Considering these results, we can infer that *C. flexuosus* EO in sublethal doses cause a similar effect as the juvenile hormone analogue, probably causing alterations in the homeostasis of hormones involved in the moulting process, which leads to abnormal larval development and growth (Dhadialla *et al*. 1998, Berti *et al*. 2013). Rattan (2010) mentioned that the EO and its constituents affect biochemical processes of insects, especially endocrine balance, with alterations in the morphogenesis process. However, this effect is not caused exclusively by the major compounds of the oil, citral, and geranyl acetate. This may be due to the synergic effect of their components, generating a greater biological response (Borrero-Landazabal *et al*. 2020).

### Measuring of juvenile hormone and moult hormone levels

With the HPLC technique coupled with mass spectrometry, we identified the signals corresponding to molt hormone (MH) and juvenile hormone III (JH III) in untreated individuals. With diflubenzuron and methoprene standards treatments, we did not detected a peak corresponding to JH III. Considering that the treatments were performed in larvae L3, it is possible that the absence of JH III is due to natural diminution of the concentration of this hormone, which is higher during the first larval stages and decreases in the L4 stage to permit the pupation and metamorphosis process (Nijhout *et al*. 2014, Hernández-Martínez *et al*. 2015). When the levels of JH III decrease at the end of the larval stage, the MH induces the change from larva to pupa and from pupa to adult (Kayukawa *et al*. 2016). Corroborating this fact, Hernández-Martínez *et al*. (2015) measured JH III changes during the gonotrophic cycle of *Ae. aegypti*, without detecting the release of methyl farmesoate, an immediate precursor of the juvenile hormone, in *Ae. aegypti* adults, but in the larval stage.

When we examined the chromatogram corresponding to larvae (L4) treated with *C. flexuosus* EO and its major compounds (citral and geranyl Acetate), there was no clear peak corresponding to MH or JH III, or any of their fractions. Considering the diminution of the JH III in L4 and the possible effect of the EO on larval development, the concentration of this hormone could have been below the limit of detection based on the calibration curves performed for the JH or MH. In relation to the MH, the alteration in the process of ecdysis and chitin formation could be altered due to the action of the majority citral component, causing changes in the concentration of this hormone.

### Conclusions

The EO from *C. flexuosus* used at sublethal concentrations has bioactivity in eggs and larvae of the *Ae. aegypti* mosquito. In the egg stage, the EO can alter the development of the embryo, decreasing the number of hatched individuals when penetrating through the chorion. In early larval stages (L1, L2, and L3), it causes an effect similar to a juvenile hormone analogue, lengthening the larval period and making pupation impossible. Additionally, the majority component citral may cause alterations in the normal process of ecdysis, a process directly related to the moulting hormone. Using HPLC technique coupled with mass spectrometry, it was possible to identify the signal corresponding to JH III and MH in untreated individuals. However, in larvae treated with *C. flexuosus* EO and its major compounds (citral and geranyl acetate), there was no clear peak. This result demonstrates an alteration in the concentration of JH III and MH hormones due to the action of EO, an effect that cannot be attributed exclusively to the major components of EO, but a synergistic effect of all its metabolites.

## Acknowledgements

We thank Prof. Dr. Elena E. Stashenko for providing the essential oils, and for their chromatographic characterisation. The authors also express their appreciation to Laboratorio espectrometria de masas of the “Universidad Industrial de Santander”, especially to the technical personnel in charge of managing the equipment.

---

**Supplementary Figure 1.**
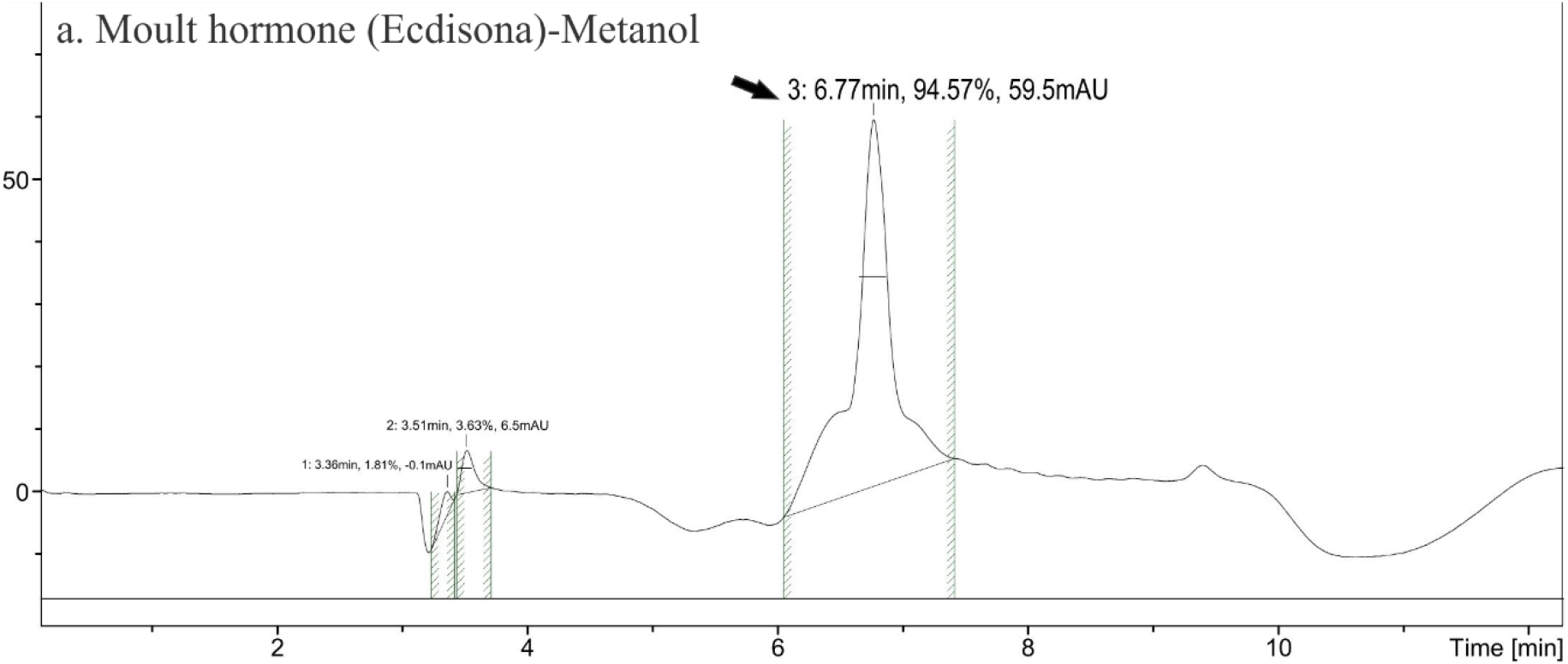
Moult hormone (MH) HPLC-mass spectrometry chromatograms of *Aedes aegypti* larvae (L4) without treatment. Y-axis: signal intensity (mAU). X-axis: retention time in minutes. Methanol-water mobile phase chromatogram. The retention time of 6.77 min, molecular mass detected by the mass spectrometer: 481.1 g.mol^-1^.

**Supplementary Figure 2.**
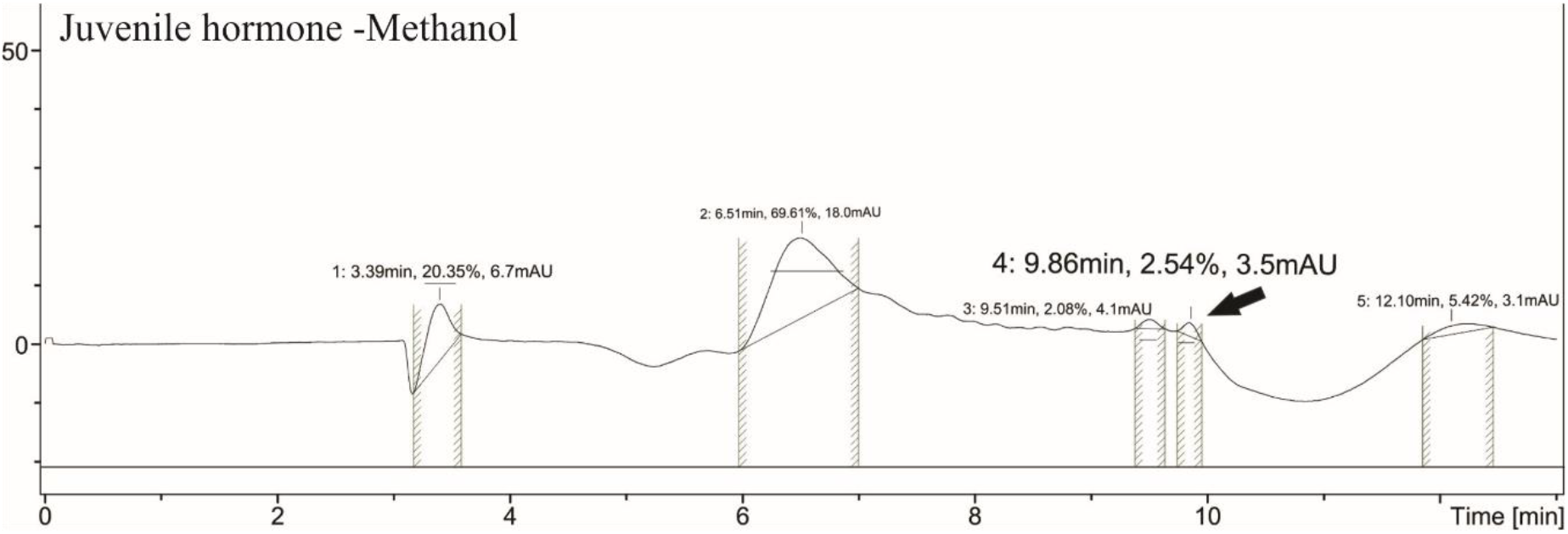
Juvenile hormone III (JH III) HPLC-mass spectrometry chromatograms of *Aedes aegypti* larvae (L4) without treatment. Y-axis: signal intensity (mAU). X-axis: retention time in minutes. Methanol-water mobile phase chromatogram. The retention time of 9.86 min.

**Supplementary Figure 3.**
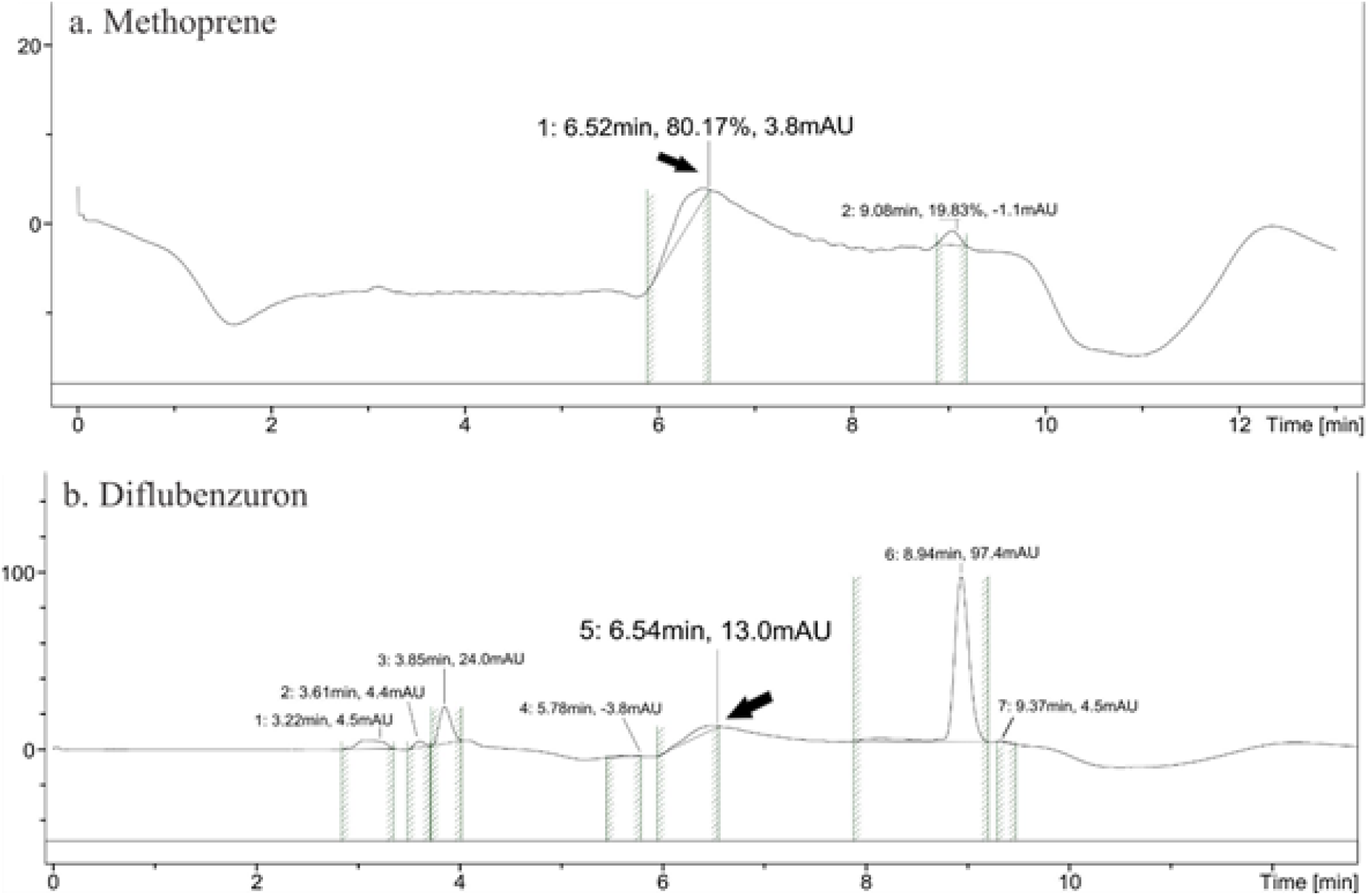
Methoprene and diflubenzuron HPLC-mass spectrometry chromatograms of *Aedes aegypti* larvae (L4). Y-axes: signal intensity (mAU). X-axes: retention time in minutes. a. *Aedes aegypti* larvae (L4) chromatogram treated with methoprene. The retention time of 6.52 min and molecular mass detected by the mass spectrometer of 481.0 g.mol^-1^ corresponds to MH. b. *Aedes aegypti* larvae (L4) chromatogram treated with diflubenzuron. The retention time of 6.54 min and molecular mass detected by the mass spectrometer of 481.1 g.mol^-1^ corresponding to MH.

**Supplementary Figure 4.**
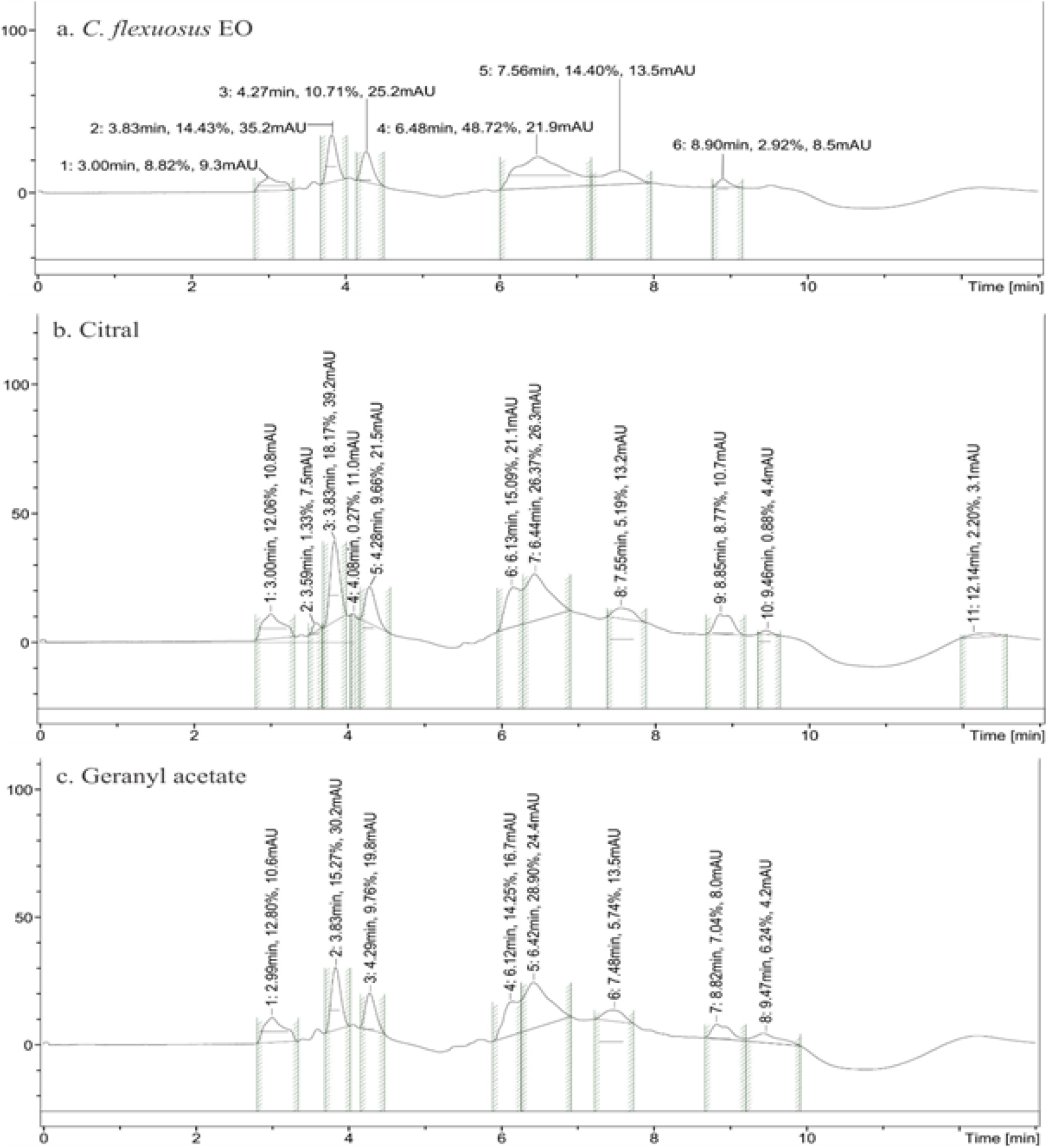
Major compounds (citral and geranyl acetate) HPLC-mass spectrometry chromatograms of *Aedes aegypti* larvae (L4) treated with *C. flexuosus* EO and its major compounds (citral and geranyl acetate). Y-axes: signal intensity (mAU). X-axes: retention time in minutes. We did not detect any titter corresponding to MH or JH III. **a.** *Aedes aegypti* larvae (L4) chromatogram treated with *C. flexuosus* EO. **b.** *Aedes aegypti* larvae (L4) chromatogram treated with the major compound citral. **c.** *Aedes aegypti* larvae (L4) chromatogram treated with the major compound geranyl acetate.

